# Supergene degeneration opposes polymorphism: The curious case of balanced lethals

**DOI:** 10.1101/2021.11.27.470204

**Authors:** Emma L. Berdan, Alexandre Blanckaert, Roger K. Butlin, Thomas Flatt, Tanja Slotte, Ben Wielstra

**Author notes:** These authors contributed equally (Emma L. Berdan, Alexandre Blanckaert).

## Abstract

Supergenes offer some of the most spectacular examples of long-term balancing selection in nature but their origin and maintenance remain a mystery. A critical aspect of supergenes is reduced recombination between arrangements. Reduced recombination protects adaptive multi-trait phenotypes, but can also lead to degeneration through mutation accumulation. Mutation accumulation can stabilize the system through the emergence of associative overdominance (AOD), destabilize the system, or lead to new evolutionary outcomes. One such outcome is the formation of balanced lethal systems, a maladaptive system where both supergene arrangements have accumulated deleterious mutations to the extent that both homozygotes are inviable, leaving only heterozygotes to reproduce. Here, we perform a simulation study to understand the conditions under which these different outcomes occur, assuming a scenario of introgression after allopatric divergence. We found that AOD aids the invasion of a new supergene arrangement and the establishment of a polymorphism. However, this polymorphism is easily destabilized by further mutation accumulation. While degradation may strengthen AOD, thereby stabilizing the supergene polymorphism, it is often asymmetric, which is the key disrupter of the quasi-equilibrium state of the polymorphism. Furthermore, mechanisms that accelerate degeneration also tend to amplify asymmetric mutation accumulation between the supergene arrangements and vice versa. As the evolution of a balanced lethal system requires symmetric degradation of both arrangements, this leaves highly restricted conditions under which such a system could evolve. We show that small population size and low dominance coefficients are critical factors, as these reduce the efficacy of selection. The dichotomy between the persistence of a polymorphism and degradation of supergene arrangements likely underlies the rarity of balanced lethal systems in nature.

## Background

Understanding the forces that maintain genetic variation despite the eroding forces of drift, directional selection, and recombination is one of the central problems of evolutionary biology. Supergenes, tightly linked sets of loci that underlie distinct complex phenotypes (1, 2), represent some of the most spectacular examples of long-term balanced polymorphisms in nature. Although the genetic architecture of supergenes does not always include chromosomal rearrangements, they are frequently present (3); here, we therefore refer to different variants of the supergene as arrangements. The reduced recombination between supergene arrangements allows for unique combinations of alleles to be maintained in linkage disequilibrium, thus preserving multi-trait phenotypes and allowing for adaptation to complex environments. However, the origin and maintenance of multiple supergene arrangements and their associated phenotypes within populations is still poorly understood (3–5).

Reduced recombination not only allows for multi-locus complexes that define supergenes, but also puts them at risk for degeneration. This reduction in recombination generates a pseudo-population substructure; effective population size (*N_e_*) for the supergene region is reduced as compared to the rest of the genome. The local decreases in *N_e_* and in effective recombination rate between supergene arrangements diminish the efficacy of purifying selection and can lead to the accumulation of deleterious alleles (3, 6, 7). Indeed, recent theoretical and empirical work has shown that supergene systems are prone to such mutation accumulation (3, 7–9). This degeneration can destabilize the system or lead to new evolutionary outcomes.

The formation of a balanced lethal system (10) has been theorized to be one such outcome (11). In a balanced lethal system, only adults possessing two distinct arrangements (i.e., heterokaryotypes) are viable. According to the rules of Mendelian inheritance, 50% of the next generation will be homozygous for either one or the other arrangement and thus inviable, meaning that half of the offspring are lost every generation. While it seems counterintuitive that such a huge genetic load could ever evolve under natural conditions, it has done so repeatedly across the tree of life, with examples in plants, insects and vertebrates (12–16). However, the degeneration of supergenes provides a potential path by which a supergene system might evolve into a balanced lethal system; a mechanism that has received very little theoretical attention (but see 17, 18). If the fitness of the homozygotes for both supergene arrangements (i.e., homokaryotypes) reaches zero, then a balanced lethal system has evolved.

Recently, Berdan and colleagues have shown that evolution of an inversion polymorphism can lead to the formation of a balanced lethal system (8). Since their focus was on the evolution of polymorphic inversions, the authors assumed that the inversion itself was inherently overdominant (*sensu stricto*), facilitating its invasion. Here, we relax this assumption, allowing only selection on linked variants. Thus in our model, associative overdominance (AOD), the heterozygote advantage experienced by a neutral variant (in this case the supergene region itself) due to selection on sites linked to the neutral locus (19–22), is the only form of balancing selection. While AOD can be caused by overdominant alleles, it can also be generated by the masking of deleterious recessive variants in multi-locus heterozygotes. Empirical and theoretical work suggests that AOD driven by deleterious recessive variants may be a strong and common force stabilizing supergene systems (6–9, 23, 24).

Here, we explore the role of AOD in maintaining a supergene polymorphism and ask when it can pave the way towards a balanced lethal system. Specifically, we examine a case where a supergene arrangement introgresses from another population. As compared to a single-population origin, the introgression scenario increases the chance of generating a balanced polymorphism, as there will be pre-existing AOD due to private deleterious recessive mutations, resulting from divergence between populations. Empirical data indicate that introgression might be a common mechanism through which supergene polymorphisms originate (25, 26). Starting from introgression of a supergene arrangement we use simulations to understand the range of conditions under which a supergene can persist and potentially evolve into a balanced lethal system.

## Model

We explored how admixture between populations, fixed for alternative arrangements of a supergene, could generate a supergene polymorphism via AOD. Following admixture, AOD is generated by the masking of recessive private deleterious mutations within the supergene region. We hypothesized that the extent of AOD and its evolutionary consequences should be based on four major factors:

1. the local mutational load of the populations,
2. the dominance coefficient of the deleterious mutations,
3. the rate of gene conversion (GC; the only recombination occurring between supergene arrangements in our model), and
4. the population size.

We simulated two isolated diploid populations (P_1_ and P_2_, *N*=2,500 individuals per population) using SLiM v2.6 (27). The genome consisted of 2 chromosomes of 10 Mb (genome length *L* = 20 Mb). We used parameters based on estimates from *Drosophila melanogaster*. Mutations occurred at a rate of *μ*=4.5 x 10^-9^ per bp per generation (28). Recombination rate was set to *r* = 4.80 x 10^-8^ per base pair per meiosis (29); recombination occurred via a combination of crossing over, at a rate of *ρ* = 3.0 x 10^−8^ per base pair per meiosis (29, 30), and gene conversion at a rate of *γ* = 1.8 x 10^−8^ per base pair per meiosis (31) (*γ* is the rate of initiation of a gene conversion event). Gene conversion track length was modeled as a Poisson distribution, with the parameter *λ* set to 500 bp (31).

To generate differences in mutational load of the populations, we used burn-ins of two different durations (*T_BI_*=200,000 or *T_BI_*=500,000 generations). Since the populations were isolated, the burn-in length corresponds to divergence time. Fitness was multiplicative, and all mutations were assumed to be deleterious. The magnitude of their fitness effects (|*s*|) was drawn from a Gamma (Γ) distribution (*α* = 0.5, *β* = 10), with fixed dominance coefficient *h*. Separate burn-ins were run for two different dominance coefficients: *h*=0 and *h*=0.1. Given that not all simulations used the same burn-in, stochasticity between burn-ins could affect possible metrics of interest when comparing across different values of *h* or burn-in length (see *<u>Significant variation with evolutionary consequences exists between burn-ins</u>* below). To take into account this source of variation, we generated 10 different burn-ins per parameter combination (2 population sizes and 2 dominance coefficients, for a total of 40 burn-ins).

We modelled the supergene as a chromosomal inversion. The inversion occurred at a fixed position on chromosome 1 and encompassed 50% of its length (i.e., 25% of the genome). The inverted arrangement was assumed to be fixed in P_2_, while the P_1_ population carried the standard arrangement. Recombination in the inverted region in the heterokaryotype was restricted to gene conversion only. The second chromosome served as a control region since indirect effects of the inversion could extend beyond the inverted region to the rest of chromosome 1.

The polymorphic period for each replicate began with the migration of a single randomly chosen individual from P_2_ to P_1_ in generation 1. This migration step established the supergene system, with the inverted arrangement from P_2_ and the standard arrangement from P_1_ forming the two arrangements of the supergene. After this migration event, we only followed evolution of P_1_. Simulations ended when the immigrating inverted arrangement was fixed or lost, or after 200,000 generations if it remained polymorphic. For each of the 10 burn-ins per parameter combination, we ran 10,000 replicates for a total of 100,000 replicates per parameter set (over 16 parameter sets in total).

We examined the emergence of AOD and its evolutionary consequences for the inversion polymorphism, using a full block design and varying two post-burn-in parameters (GC and population size) for each of the four combinations of the burn-in parameters (see Table 1). The absence of GC was modelled as having all recombination events occur as crossing overs, so that the total rate of recombination, *r*, remained constant. To explore the effect of a smaller population size, we assumed that a bottleneck occurred before the migration event, with only 100 individuals among the 2,500 in P_1_ contributing to generation 1 of the AOD period. The polymorphic period differed from the burn-in period in that beneficial mutations could also occur at a 1,000x lower rate than their deleterious counterparts (4.5 x 10^-12^ per bp per generation). All beneficial mutations were co-dominant (*h*=0.5) and their fitness effects (|*s*|) were drawn from an exponential distribution with parameter κ = 0.001.

**Table 1.** Parameters for the simulations

| Parameter | Value |
| --- | --- |
| Mutation rate ($\mu$) | $4.5 \times 10^{-9}$ per bp per generation |
| Genome length ($L$) | 20 Mb |
| Recombination ($r$) | $4.80 \times 10^{-8}$ per bp per meiosis |
| Gene conversion initialization ($\nu$) | 0 or $1.8 \times 10^{-8}$ per base pair per meiosis |
| Gene conversion tract length ($\lambda$) | 500 bp |
| Burn-in length ($T_{BI}$) | 200,000 or 500,000 generations |
| Dominance coefficient ($h$) | 0 or 0.1 |
| Population size post-burn-in ($N$) | 100 or 2,500 (250 and 500 for selected combinations of other parameters) |

To further explore the relationship between population size and the emergence of balanced lethals, we considered two additional population sizes (*N*=250 and *N*=500; resulting from a bottleneck of the *N*=2,500 population, as described above) and tested these with the two different burn-in lengths. Both GC and *h* were kept at 0, as these factors were found to be critical for the emergence of balanced lethals using our initial parameter combinations (see *<u>Balanced lethals can only evolve in a highly restricted parameter space</u>* below). As before, we performed 10,000 replicate runs of each of these parameter sets, for each of the 10 burn-ins, for a total of 100,000 replicates.

To reduce computational time we did not simulate neutral mutations. We simulated allelic content only in 200 kb of the 20Mb genome, divided into 5 kb segments, uniformly distributed along the genome. Recombination can occur anywhere in the genome, but deleterious and beneficial mutations only occur in regions where we simulated allelic content. Finally, we scaled our parameters by a factor of 10, keeping *Lμ* and *Lr* constant, so that evolution occurred at an accelerated rate (a common practice; see (32) for an example).

Since we assumed a constant population size, only relative fitness values were considered. For the rest of the paper, we refer to fitness relative to the mean fitness of the population.

## Results and Discussion

### Significant variation with evolutionary consequences exists among burn-ins

The mutational load of the populations, following the burn-in, is affected by dominance and burn-in length. As expected, the number of fixed mutations between different populations increases with burn-in length, but the number of segregating mutations remains stable (figure S1). Mutations accumulate in a linear fashion, with the 500,000 generations burn-in having ~2.5x more fixed mutations than the 200,000 generation burn-in. Increasing the dominance coefficient (*h*) strongly decreases the number of segregating mutations (figure S1B). Additionally, increasing the dominance coefficient slightly decreases the average number of fixed mutations, however the distributions overlap (figure S1A). Thus, by simply picking randomly a single burn in for each value of *h*, there is a non-negligible chance that the average trend observed would be reversed, highlighting the necessity of using multiple burn-ins. These effects are mirrored for mutational load (figure S2). The mutational load of the supergene region, especially in P_1_ (the focal population), is strongly correlated with the strength of AOD at the beginning of the simulation (figure 1A, figure S3). We calculate AOD following Ohta (19), using *s’* = min(*s’_1_,s’_2_*), where:

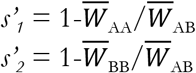

**Figure 1.**
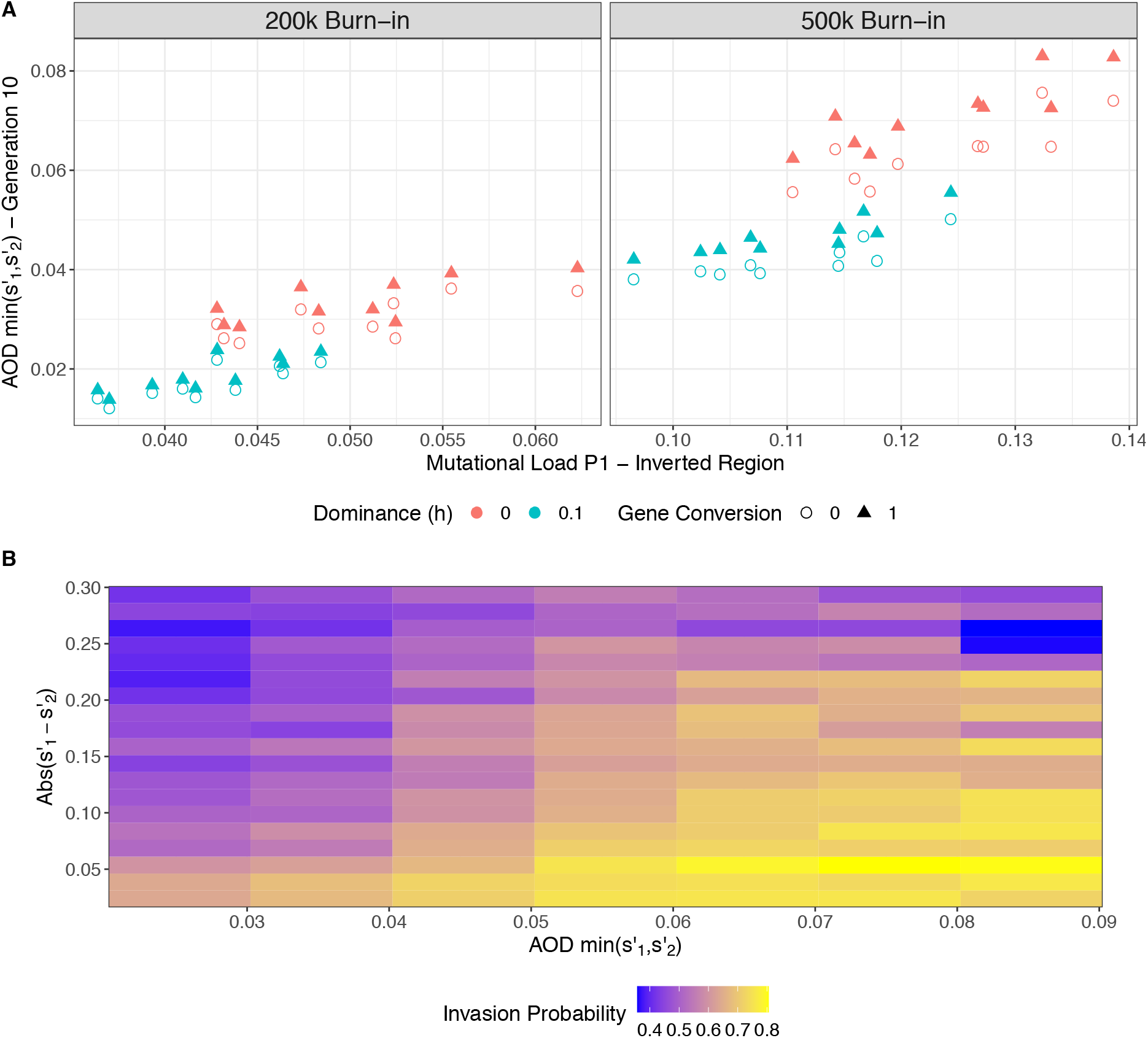
Associative overdominance (AOD) generated from mutational load can facilitate invasion of a supergene. A.) AOD correlates with mutational load. Shown is the average AOD at generation 10, across replicates, for each burn-in (200k or 500k generations), compared to the mutational load of the P_1_ population estimated from the burn-in. Colors indicate the dominance coefficient (0-red, 0.1-blue) and shape indicates the level of gene conversion (circle - absence, triangle - presence). The data shown is for *N*=2,500. B.) Both symmetry and strength of AOD contribute to the probability of invasion. Each tile represents a specific parameter space (+/-0.005 change in AOD and +/-0.0075 change in absolute (*s*’_1_-*s*’_2_)). Coloring represents the probility that a simulation from that parameter space resulted in invasion. The number of obervations per tile differs, but all are greater than 20. The data shown is for *N*=100, GC=0, and *h*=0.

Where A is the inverted arrangement (originating in P_2_) and B is the standard arrangement (originating in P_1_). Positive values of *s’* indicate that the heterokaryotype is more fit than either homokaryotype (i.e., the presence of AOD) and negative values indicate the reverse. Variation in mutational load and in the strength of AOD is found across parameter combinations, as well as among burn-ins within parameter combinations (figure 1A).

The strength of AOD at the beginning of the simulation partially explains the probability of invasion (figure 1B). We consider that the inverted arrangement has successfully invaded if it is still present in P_1_ after *N* generations. As AOD increases, so does the probability of invasion. This is in line with theoretical predictions that AOD should aid supergene invasion (23).

### AOD is evolutionarily unstable because it evolves

AOD is determined by the mutational loads of the two supergene arrangements. Its strength therefore varies as the supergene arrangements evolve. Nei *et al*. found that AOD could aid the invasion as well as the fixation of a new inversion (33). Yet, in many circumstances such an advantage conferred by AOD might only be transient as the inversion will accumulate recessive deleterious mutations over time which might in turn prevent the fixation of the inversion homokaryotype (34). In our model, where the inversion is introduced by migration, invasion probability is also dependent on the symmetry of the mutational load of the two supergene arrangements in addition to the strength of AOD (figure 1B). This is consistent with results from single-locus overdominance models which show that higher levels of heterozygosity can be maintained under symmetrical overdominance, because drift is less likely to cause the loss of a polymorphism maintained at intermediate frequency (35, 36). A critical difference between AOD and single-locus overdominance is that AOD can easily evolve (34), while the overdominance coefficient is generally a fixed property of the mutation.

AOD is determined by large regions of the genome that are prone to mutation accumulation due to reduced recombination (8, 9). The rate of mutation accumulation in either of the supergene arrangements is tied to both effective recombination rate and the relative strengths of drift and purifying selection (correlated with the number of copies of the arrangement). A large difference between the fitness of the supergene arrangements implies a large frequency difference between the major (more frequent) and minor (less frequent) supergene arrangements. This in turn translates into purifying selection being far less effective in the minor arrangement, which leads to an increase in mutation accumulation (8). This has two consequences: first the relative fitness of the major arrangement will increase even more, further increasing its frequency, and therefore the efficacy of purifying selection, in a feedback loop. Second, if mutations are only partly recessive, mutation accumulation in the minor type will also decrease the fitness of the heterokaryotype and thus reduce AOD. These processes drive the frequency of the minor arrangement down, increasing its chance of being lost by drift. To illustrate this point, consider a single-locus model with overdominance and assume that one homozygote is inviable (see Appendix). In order for the frequency of the corresponding allele to remain above 10%,

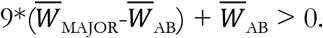

Using our notation, this can be simplified (regardless of dominance) to:

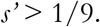

The critical factors for the maintenance of the supergene polymorphism are the fitness differential between the major arrangement and heterokaryotype and the mean fitness of the heterokaryotype. An increase in dominance coefficient will reduce both terms; as predicted, we observe a negative relationship between dominance coefficient and invasion probability (figure S4). Together, our results point towards the symmetry of the mutational load of the two supergene arrangements (along with the magnitude of AOD) as being a key factor determining the fate of the supergene polymorphism.

AOD readily evolves in our model: it generally decreases in the beginning of the runs and then increases again over time (figure 2). This remains true even when focusing only on simulations where the population remains polymorphic to the end. This indicates that this increase is not due to a sieving effect (i.e., cases with weak AOD are removed at early time points leaving only cases with strong AOD at later time points). In addition, the initial decrease in AOD occurs even in the absence of gene conversion and with fully recessive deleterious mutations, excluding the possibility that the mutational load of the immigrating arrangement decreases via gene flux and selection. Thus, this initial decrease is likely due to the erosion of linkage disequilibrium between the supergene arrangement from P_2_ and the rest of the P_2_ genome. Initially, the entire chromosome 1 (50% of the genome) contributes to overdominance, but over time recombination will reduce this to the supergene alone. Indeed, this drop in AOD corresponds to a drop in heterokaryotype fitness and a slight increase in the fitness of the inversion homokaryotype (figure S5). After the initial decrease, AOD increases in all runs that remain polymorphic over 200,000 generations, due to mutation accumulation. This increase is stronger in the absence of GC, with fully recessive deleterious mutations, and in smaller populations (figure S6).

**Figure 2.**
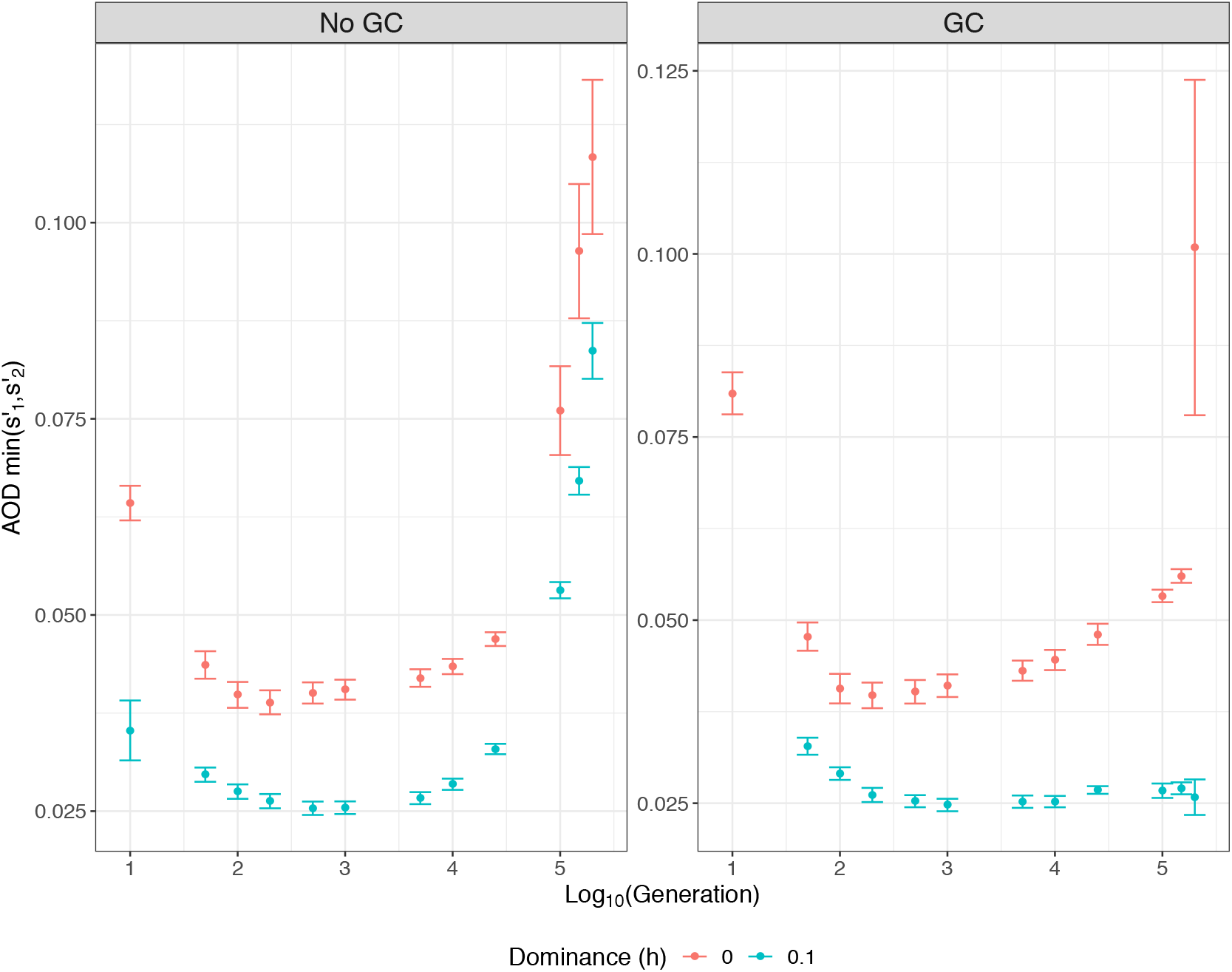
Associative Overdominance (AOD) evolves in a non-monotonous manner over time. Shown are changes in AOD over time, only for simulations where the polymorphism lasted 200,000 generations (*N*=2,500, Burn-in=500k generations). AOD for each generation is calculated as an average per burn-in, per parameter set. Error bars represent standard error between burn-ins. Colors indicate the dominance coefficient (0-red, 0.1-blue) and facets indicate the level of gene conversion (GC; presence or absence)).

### Maintenance of supergene polymorphism can be accomplished in three different ways

To understand the fate of the supergene and its possible long-term maintenance, we focus on cases where the polymorphism is maintained over the full simulation period (i.e., 200,000 generations). We examine fitness across karyotypes at the final generation, considering individuals with fitness values < 0.01 to be inviable and individuals with fitness values > 0.01 to be viable. We identify three qualitatively different outcomes:

1. both homokaryotypes remain viable (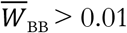 & 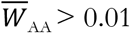),
2. one homokaryotype becomes inviable, while the other remains viable (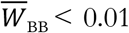 & 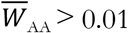 or 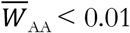 & 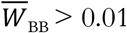), and
3. both homokaryotypes become inviable and only heterokarytoypes contribute to subsequent generations (i.e., a balanced lethal system; 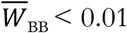 & 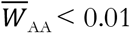).

Outcome 1 only occurs in the large populations (*N*=2,500) and is negatively correlated with GC and positively correlated with *h* and divergence time before introgression (figure S7). Outcome 2 occurs almost exclusively in large populations (*N*=2,500; 99.99% of cases), without GC (99.92% of cases), and is negatively correlated with *h* and positively correlated with divergence time (figure S8A). Finally, outcome 3 (a balanced lethal system) only occurs in the small populations (*N*=100) and is negatively correlated with *h* and positively correlated with divergence time (figure S8B). Overall, either partially recessive mutations or GC, or both, reduce the chance of the supergene remaining polymorphic for up to 200,000 generations (i.e. at least 80 *N* generations).

We further explore how fitnesses change over time when the polymorphism persists. When both homokaryotypes remain viable (outcome 1), there is an initial increase in fitness of all three karyotypes (figure 3). In our simulations there are three ways for a homokaryotype to increase in fitness: (i) purging of deleterious mutations via gene flux, (ii) reducing linkage disequilibrium with the rest of the genome, therefore lessening the mutational load generated by private mutations, and (iii) accumulation of beneficial mutations. Recombination can occur within arrangements via crossing over and GC or between arrangements via GC only. The introgressed inverted arrangement cannot purge its initial mutational load by recombination within arrangements, since all inverted arrangements share the same load. We observe increases in fitness, both in the presence and absence of GC. This indicates either accumulation of beneficial mutations or the lessening of linked mutational load generated by private mutations (figure 3). After the initial increase, fitness of the homokaryotypes plateaus, before decreasing again. This decrease is due to the accumulation of deleterious mutations, which do not affect the fitness of the heterokaryotype when *h*=0. It is important to remember that we only run our simulations for 200,000 generations. Given that karyotype fitnesses are still evolving, outcome 1 is likely a transient state, as *s’* is quite small and asymmetric mutation accumulation could result in the loss of the polymorphism.

**Figure 3.**
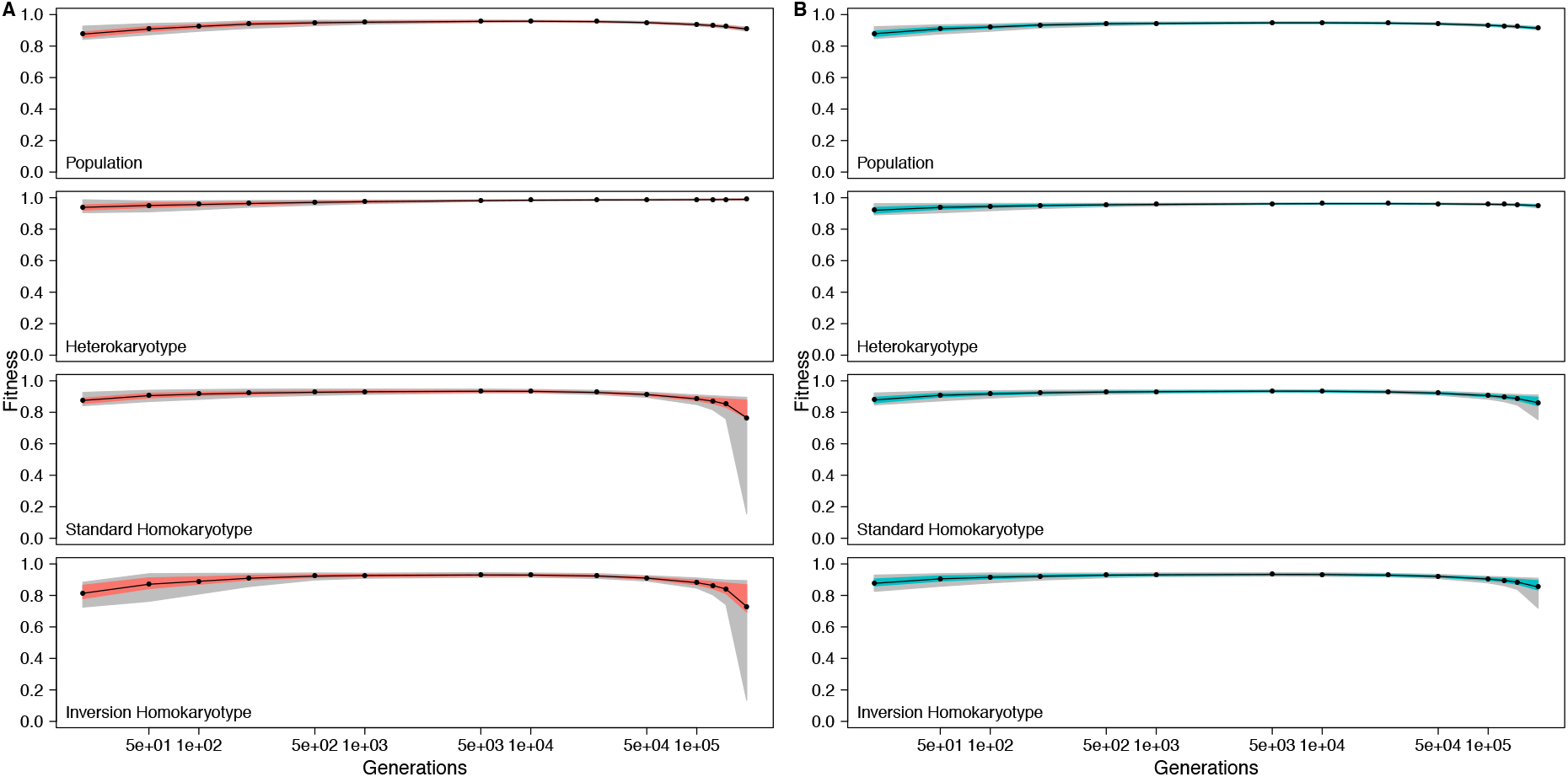
Evolution of fitness when all karyotypes remain viable for different values of *h* (A. *h*=0, B. *h*=0.1). Fitness of the population, relative fitness of the heterokaryotype, relative fitness of the standard homokaryotype, and relative fitness of the inversion homokaryotype are shown. Time is always shown on a log scale, actual time points are: 10, 50, 100, 200, 500, 1000, 5000, 10000, 25000, 50000, 100000, 125000, 150000, and 200000 generations. Only data from *N*=2,500 runs that lasted 200,000 generations are shown.

In the half-lethal outcome (outcome 2), one arrangement degrades by mutation accumulation, while the other does not (figure S9). Initially, fitnesses of all karyotypes increase, due to the accumulation of beneficial mutations and the reduction of linkage disequilibrium between the supergene and the other P_2_ mutations situated outside of the supergene. After reaching a plateau, the fitness of one of the homokaryotypes drops steeply to “lethal” (fitness < 0.01), due to the accumulation of deleterious mutations, while the other remains relatively fit (figure S9). The fitter arrangement also accumulates mutations over time, but at a much slower rate (figure S9). Outcome 2 is likely to be less transient than outcome 1 (where both homokaryotypes remain viable), because there is the possibility for *s’* to increase as the fitter homokaryotype slowly degrades. This indicates that AOD is able to maintain a long-term polymorphism when at least one of the homokaryotypes is inviable. Indeed, many supergene systems have a single lethal homokaroytype, including e.g. the ruff (37, 38) and the fire ant (39, 40). We discuss outcome 3 below (see section *<u>Balanced lethals can only evolve in a highly restricted parameter space</u>*).

### The degradation of supergene arrangements and the maintenance of the polymorphism via AOD are at odds

The parameter space within which the supergene polymorphism might potentially be maintained over long time-scales by AOD alone is small. Events occurring with a frequency below 10^-4^ were ignored. Outcomes 1 and 2 only occur in 5 and 2 parameter combinations, respectively, out of 16. This suggests, that within our model, AOD alone is only capable of maintaining a long-term balanced polymorphism in exceptional circumstances. We hypothesize that this is due to the fact that degeneration of the supergene arrangements, the process that increases AOD, typically also leads to homokaryotype asymmetry.

To test this idea, we examine how our four investigated parameters (GC, dominance coefficient, divergence time, and population size) affect critical properties (Table 2). For a polymorphism to be maintained, the inverted arrangement must invade, but not fix. This creates a challenge; the underlying cause of the necessary fitness advantage of the heterokaryotype, should not also generate a fitness advantage for the inverted homokaryotype. For instance, increased divergence between populations increases both invasion and fixation probability. Stronger initial AOD means that the immigrating arrangement quickly spreads in the population, thus spending less time at low frequency, where mutation accumulation is more likely. Once this happens, and if the two arrangements are approximately equivalent in load, it is unpredictable which one is fixed or lost.

**Table 2.**
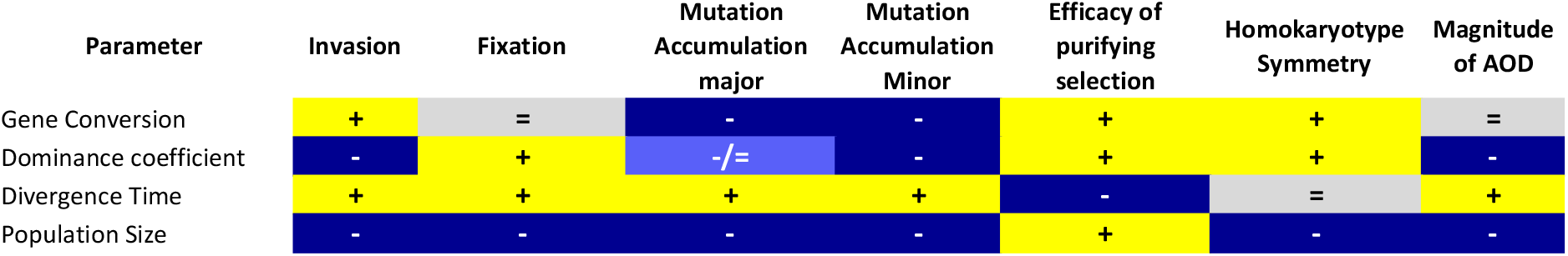
Shown are the relationship between the parameter and process (yellow positive, blue negative, grey no relationship). Note that dominance coefficient refers to the coefficient for deleterious mutations only.

If invasion succeeds, the processes that will decide the fate of the polymorphism are the rates of mutation accumulation in the major and minor arrangements. Only mutation accumulation in the major arrangement will increase *s’* and symmetry. Mutation accumulation in the minor arrangement will not affect *s’* but will decrease symmetry. Due to the feedback loop between the effective population size of each supergene arrangement and mutation accumulation (discussed above in *<u>AOD is evolutionarily unstable because it evolves</u>*), mutation accumulation is (much) faster in the minor arrangement. The presence of the feedback loop thus makes maintenance of the polymorphism by AOD alone unlikely.

Several factors impact mutation accumulation in the major and minor arrangements and the resulting fate of the polymorphism (figure 4). First, GC, by allowing recombination between the two arrangements, can reduce asymmetry, promoting the polymorphism. Surprisingly, we do not find a detectable reduction of the magnitude of AOD associated with GC. This might be due to the low population-wide GC rate (*N_γ_*) relative to the rate of mutation accumulation. Second, incompletely recessive deleterious mutations are partially expressed in the heterokaryotype. This results in lower AOD, making the polymorphism more likely to be lost. However, the partial expression in the heterokaryotype also allows purifying selection to act on the mutations, especially those in the minor arrangement. This slows the rate of mutation accumulation and the build-up of the asymmetry (figure 4). Third, population size has an unexpected role here. Small populations mean that maintenance of the polymorphism becomes unlikely, as drift becomes stronger, with the exception of one case: the balanced lethal system, which we discuss below. On the other hand, larger population size means that selection can act upon smaller fitness differences. With stronger purifying selection, a tiny initial asymmetry in mutation load can trigger a ‘snowball effect’, leading to the loss of one of the two arrangements, even if they start off in perfect symmetry. Finally, the divergence time between the two populations may be the only factor that helps to maintain the polymorphism following invasion. A stronger initial mutational load in both arrangements means a stronger initial AOD plus lower asymmetry between arrangements because of the larger sample of mutations.

**Figure 4.**
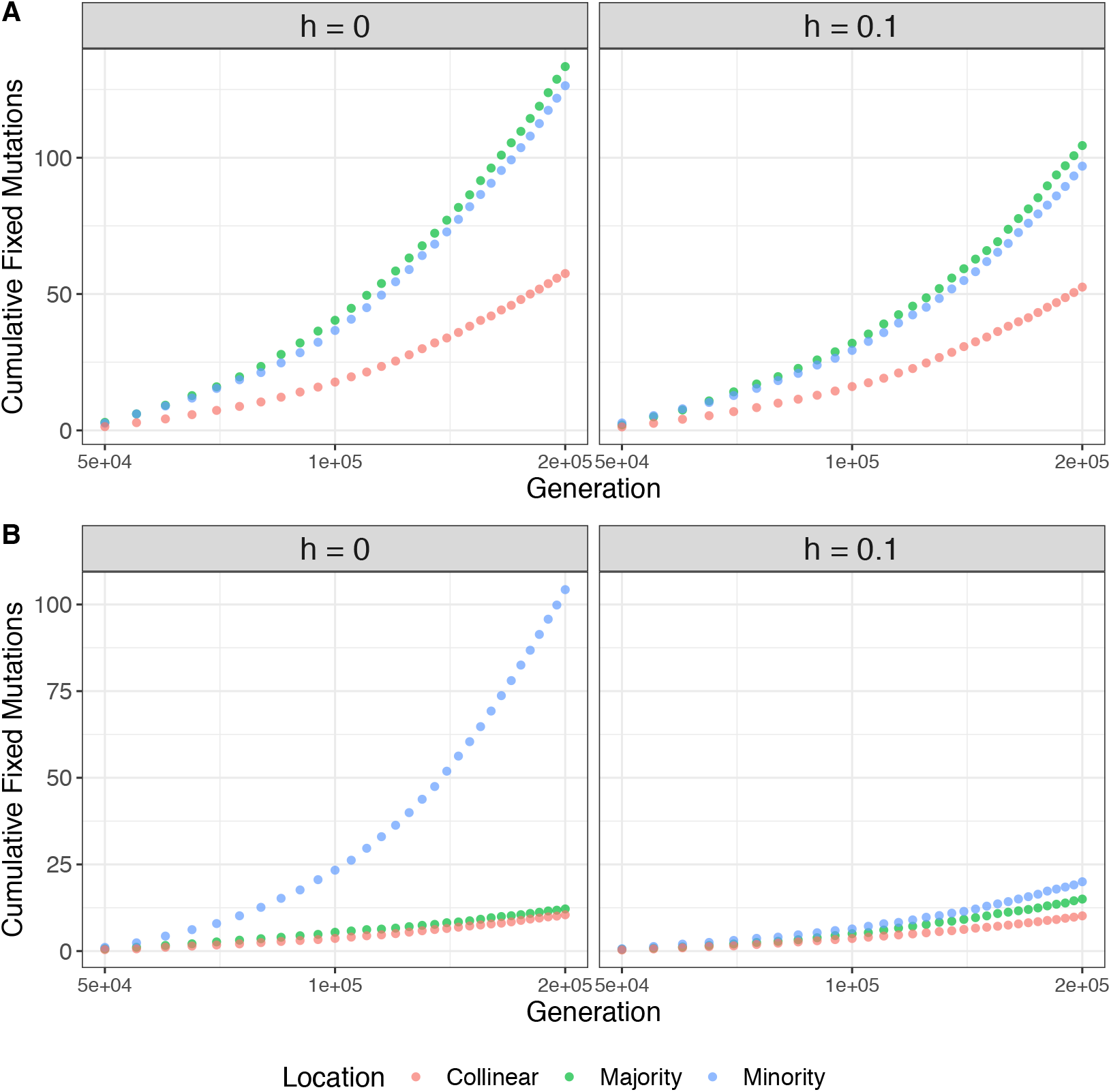
Mutation accumulation varies across parameter space. Shown is cumulative accumulation of mutations in (A) small populations (*N*=100) and (B) large populations (*N*=2,500) in the major arrangement (green), the minor arrangement (blue), and the unlinked collinear region (red). Facets indicate the dominance coefficient of deleterious mutations. For all data there is no gene conversion and only runs that lasted 200,000 generations are shown.

Up to this point we have only considered balancing selection caused by AOD driven by deleterious recessive mutations. However, other forms of balancing selection may help to stabilize the polymorphism. Negative frequency-dependent, spatially and/or temporally varying selection are well-known forms of balancing selection that are found in supergene systems (1, 3, 5, 41). One particular mechanism, disassortative mating, is predicted to evolve in systems with strong heterozygote advantage (e.g., AOD 42). Indeed, there is evidence of disassortative mating in many classic supergene systems that also show signs of mutation accumulation (25, 26, 43–45).

### Balanced lethals can only evolve in a highly restricted parameter space

The evolution of a supergene system into a balanced lethal system only happens under a stringent set of conditions. We only observe the evolution of balanced lethals in small populations. The probability of a balanced lethal outcome decreases with population size and is not observed for N>=500. (figure 5). This is because the feedback loop between arrangement frequency and mutation accumulation is disrupted by drift in small populations. Specifically, purifying selection is going to be less effective in general, and in the major arrangement in particular. Additionally, arrangement frequencies will fluctuate more between generations due to drift. Together, these hamper the snowballing of an initial asymmetry by selection, occasionally allowing for both supergene arrangements to degrade simultaneously (figure 5). However, in a small population, loss of the polymorphism is the most common outcome, meaning that balanced lethal systems remain rare.

**Figure 5.**
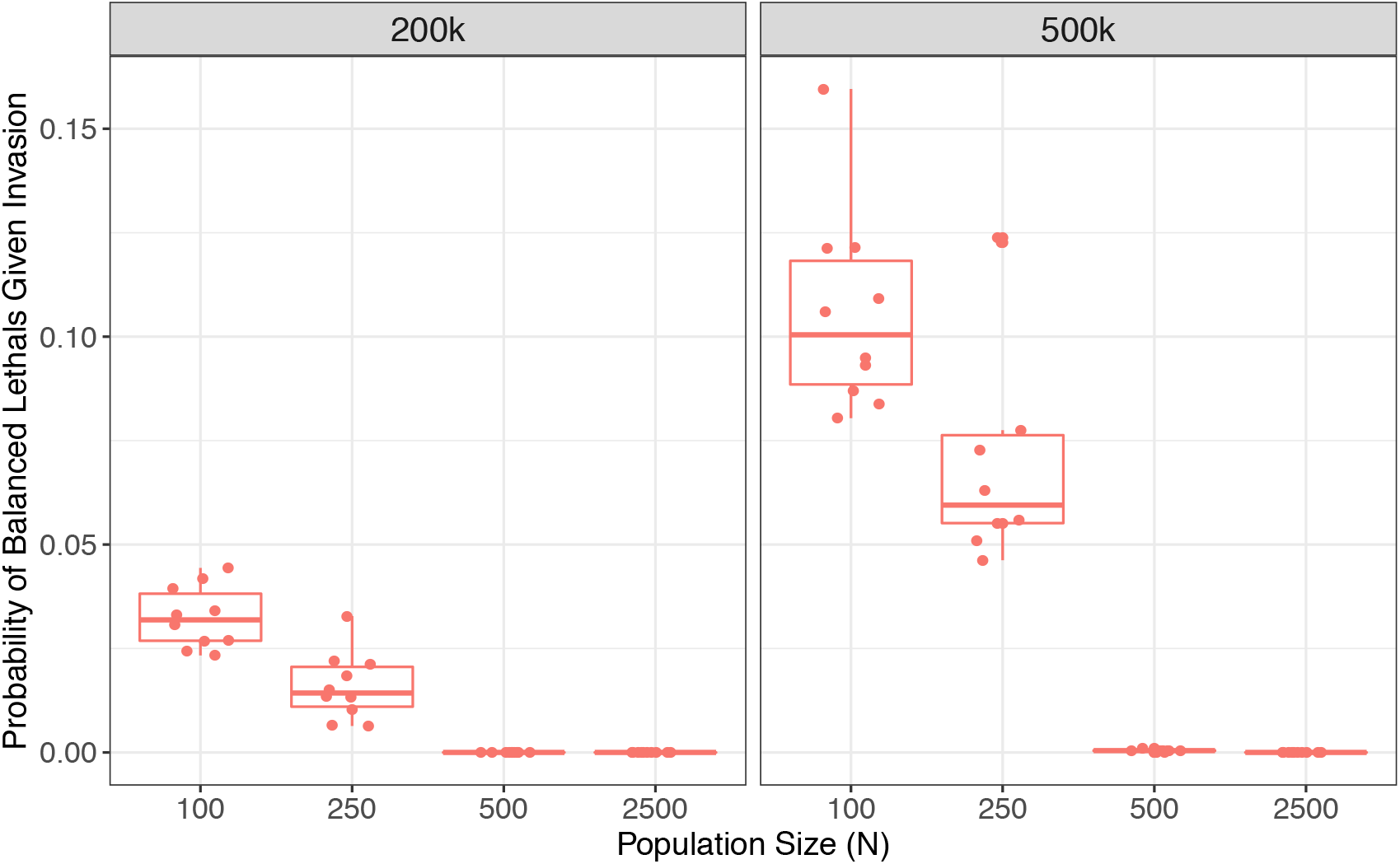
Balanced lethals evolve under restricted conditions. Probability of balanced lethals given invasion is shown as a function of the population size in the absence of gene conversion and for fully recessive mutations (*h*=0). Facets indicate burn-in length (200k or 500k generations). Each point is a burn-in.

Another crucial factor is the dominance coefficient of the deleterious mutations. While we observe the evolution of balanced lethals when mutations are not fully recessive, the likelihood of observing this outcome is reduced. This is because partially recessive mutations are expressed in the heterokaryotype, reducing AOD and destabilizing the polymorphism. In contrast, the presence of GC does not seem to affect the outcome in line with the fact that it has little impact on AOD overall.

## Conclusions

While AOD can lead to the establishment of a supergene system, it is unlikely to maintain polymorphism over long time periods when acting alone. Supergenes degenerate over time as the decrease in effective population size and recombination rate leads to mutation accumulation. Initial asymmetries often mean that one of the two supergene arrangements degenerates more rapidly, which makes polymorphism sensitive to loss through drift.

We observe only two potentially persistent polymorphic states in our model, a half-lethal system and a fully balanced lethal system, both of which occur only in small parts of parameter space. Importantly, both states are found in natural supergene systems (3, 11). As AOD alone is unlikely to maintain the supergene over long time scales, other forms of balancing selection, or disassortative mating, might be key to maintaining supergene polymorphisms. This matches empirical evidence from multiple supergene systems, where complex multi-dimensional selection is involved in the maintenance of polymorphism (3, 4, 43, 46, 47).

In our simulations, balanced lethals only occur in small populations under restrictive conditions. This finding is in good qualitative agreement with the fact that, despite a broad representation across the tree of life, balanced lethal systems appear to be quite rare (10). We anticipate that advances in genomics will provide insights into the frequency with which supergenes degrade and evolve into balanced lethal systems.

## Methods

The simulation code and analysis code is available at gitlab.com/slottelab/BL_sims

## Acknowledgements

We thank Inês Fragata and Brian Charlesworth for helpful discussions.

## Supplemental Figure Legends

Figure S1. Differences in fixed and segregating mutations between burn-ins. A.) Number of fixed mutations between P1 and P2, B.) Number of segregating mutations in P2. Colors indicate the dominance coefficient (0-red, 0.1-blue).

Figure S2. Differences in mutational load between burn-ins. A.) Mutational load of P1 B.) Mutational load of P2. Colors indicate the dominance coefficient (0-red, 0.1-blue).

Figure S3. Mutational load of P2 correlates with associative overdominance (AOD). Facets indicate burn-in length, shape indicates rate of gene conversion (absent or present) and color indicates the dominance coefficient (0-red, 0.1-blue). The data shown is for *N*=2,500.

Figure S4. Dominance decreases invasion probability. Probability of invasion is shown for all parameter sets. Facets indicate *N* (100 or 2,500), and gene conversion (GC; absent or present). Colors indicate the dominance coefficient (0-red, 0.1-blue). Each point is a different burn-in.

Figure S5. Changes in fitness within the first 1,000 generations after invasion. Shown are normalized fitnesses for A.) the inversion homokaryotype, B.) the standard homokaryotype, C.) the heterokaryotyope, and D.) the mean fitness of the entire population. Normalized fitness for each generation is calculated as an average per burn-in per parameter set. Error bars represent standard error between burn-ins. Facets indicate gene conversion (GC; absent or present) and colors indicate the dominance coefficient (0-red, 0.1-blue). *N*=2,500 and burn-in length = 500,000 for all graphs.

Figure S6. The evolution of associative overdominance (AOD) under different parameters. Shown is the evolution of AOD over time for simulations where the polymorphism lasted 200,000 generations A.) *N*=100, and B.) *N*=2,500. AOD for each generation was calculated as an average per burn-in per parameter set. Error bars represent standard error between burn-ins. Colors indicate the dominance coefficient (0-red, 0.1-blue) and facets indicate the level of gene conversion (absent or present). Burn-in length=500,000 for all graphs. Note the change in y-axis scale.

Figure S7. Polymorphism of the inversion with fit homokaryotypes given invasion. Probability that the supergene is polymorphic at the end of the simulation and both homokarotypes are fit is shown for all parameter sets. acets indicate *N* (100 or 2,500), and gene conversion (GC; absent or present). Colors indicate the dominance coefficient (*h*; 0-red, 0.1-blue). Each point represents a different burn-in.

Figure S8. A. Probability of a half lethal system given invasion. Probability that the supergene is polymorphic at the end of the simulation and one homokarotypes is lethal is shown for all parameter sets. B. Balanced lethal outcome given invasion. Probability that the supergene has eveoved into a balanced lethal system after invasion. For both plots facets indicate *N* (100 or 2,500), and gene conversion (GC; absent or present). Colors indicate the dominance coefficient (*h*; 0-red, 0.1-blue). Each point represents a different burn-in.

Figure S9. Evolution of karyotype fitnesses in a half lethal system. Fitness of the population, relative fitness of the heterokaryotype, relative fitness of the inversion homokaryotype, and relative fitness of the standard homokaryotype are shown for cases where (A) The inversion homokaryotype becomes lethal, (B) The standard homokarotype becomes lethal. Time is always shown on a log scale, actual time points are: 10, 50, 100, 200, 500, 1000, 5000, 10000, 25000, 50000, 100000, 125000, 150000, and 200000 generations. For all data shown *N*=2,500, gene conversion is absent and all mutations are fully recessive (*h*=0).

