## Supplemental Figures for "Supergene degeneration opposes polymorphism: The curious case of balanced lethals"

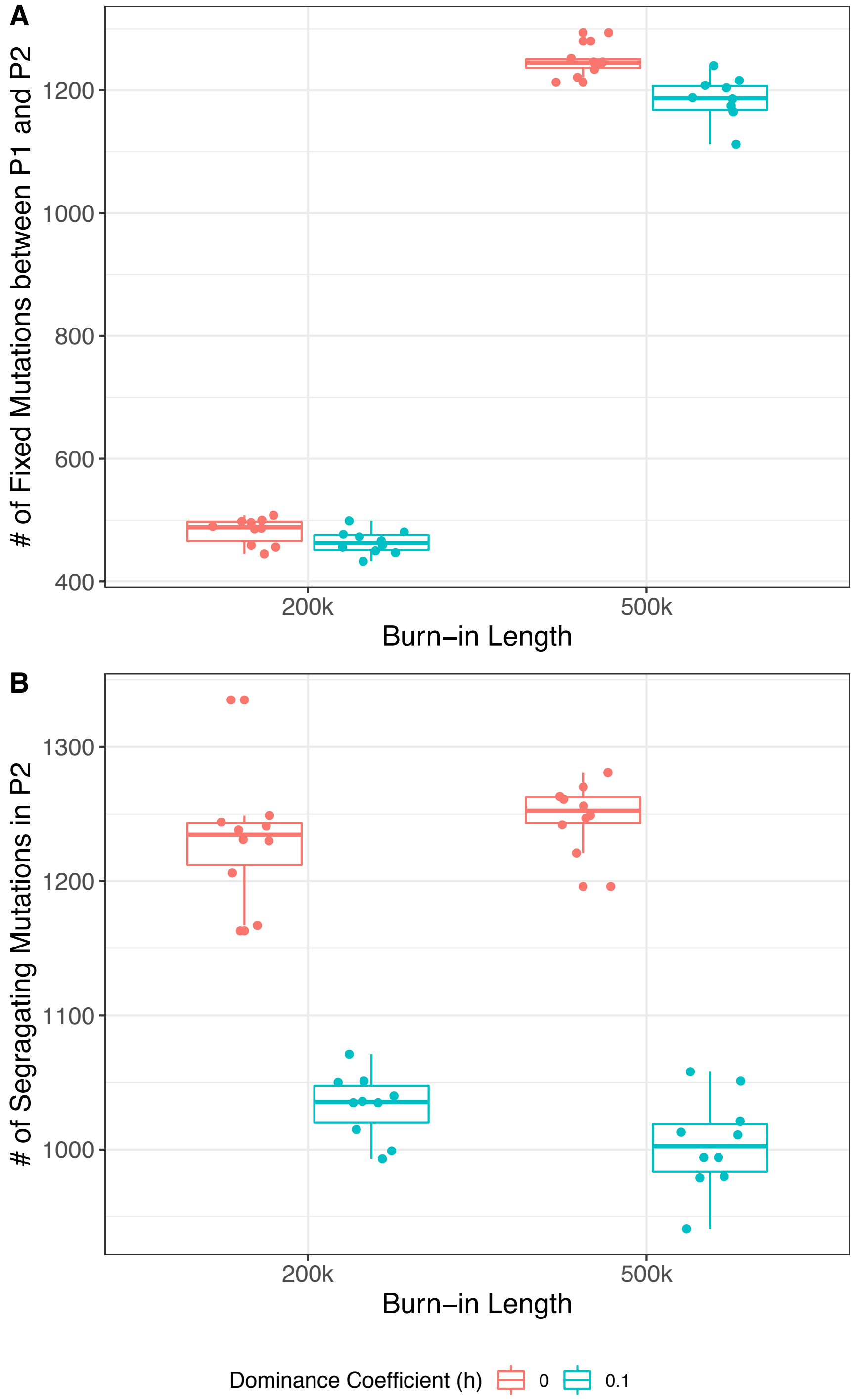


Figure S1. Differences in fixed and segregating mutations between burn-ins. A.) Number of fixed mutations between P1 and P2, B.) Number of segregating mutations in P2. Colors indicate the dominance coefficient (0-red, 0.1-blue).

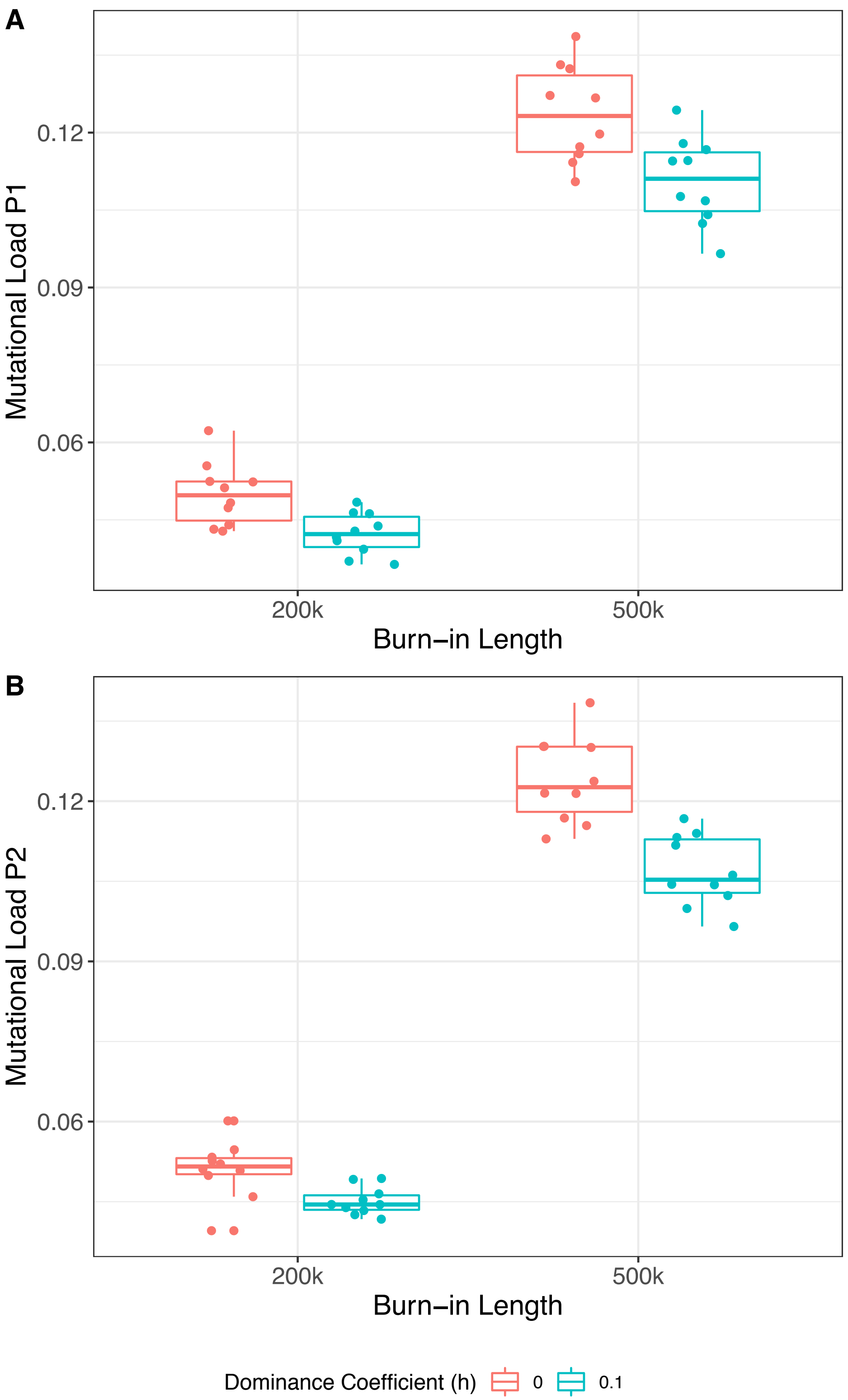


Figure S2. Differences in mutational load between burn-ins. A.) Mutational load of P1 B.) Mutational load of P2. Colors indicate the dominance coefficient (0-red, 0.1-blue).

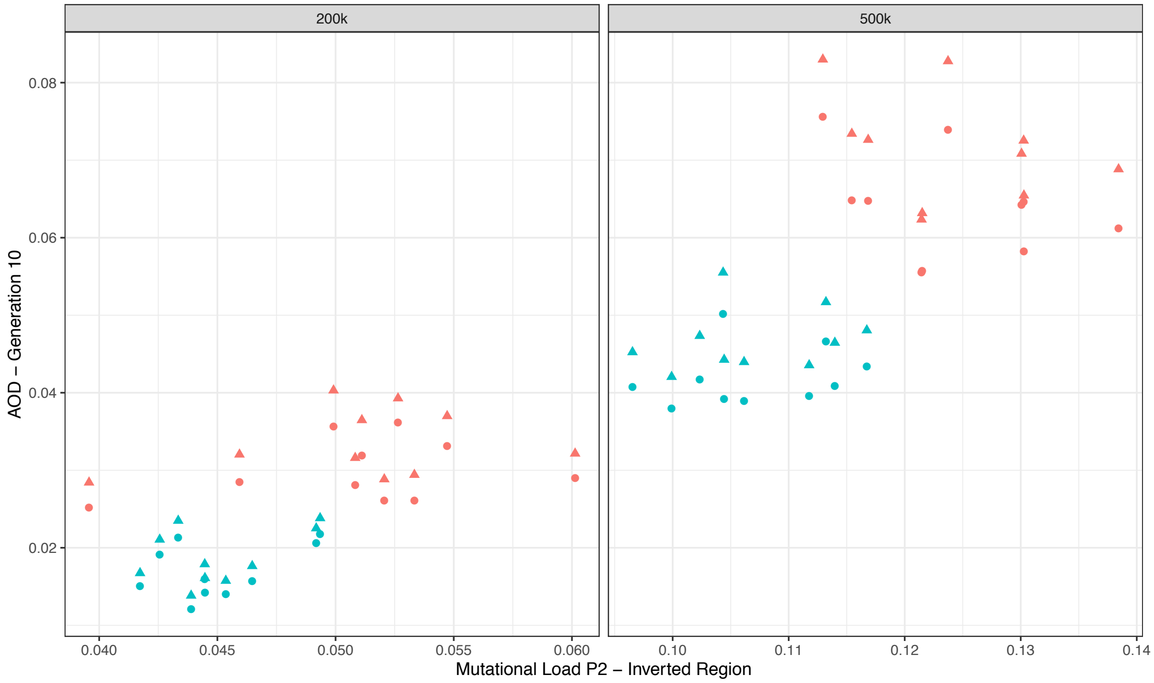


Figure S3. Mutational load of P2 correlates with associative overdominance (AOD). Facets indicate burn-in length, shape indicates rate of gene conversion (absent or present) and color indicates the dominance coefficient (0-red, 0.1-blue). The data shown is for N=2,500.

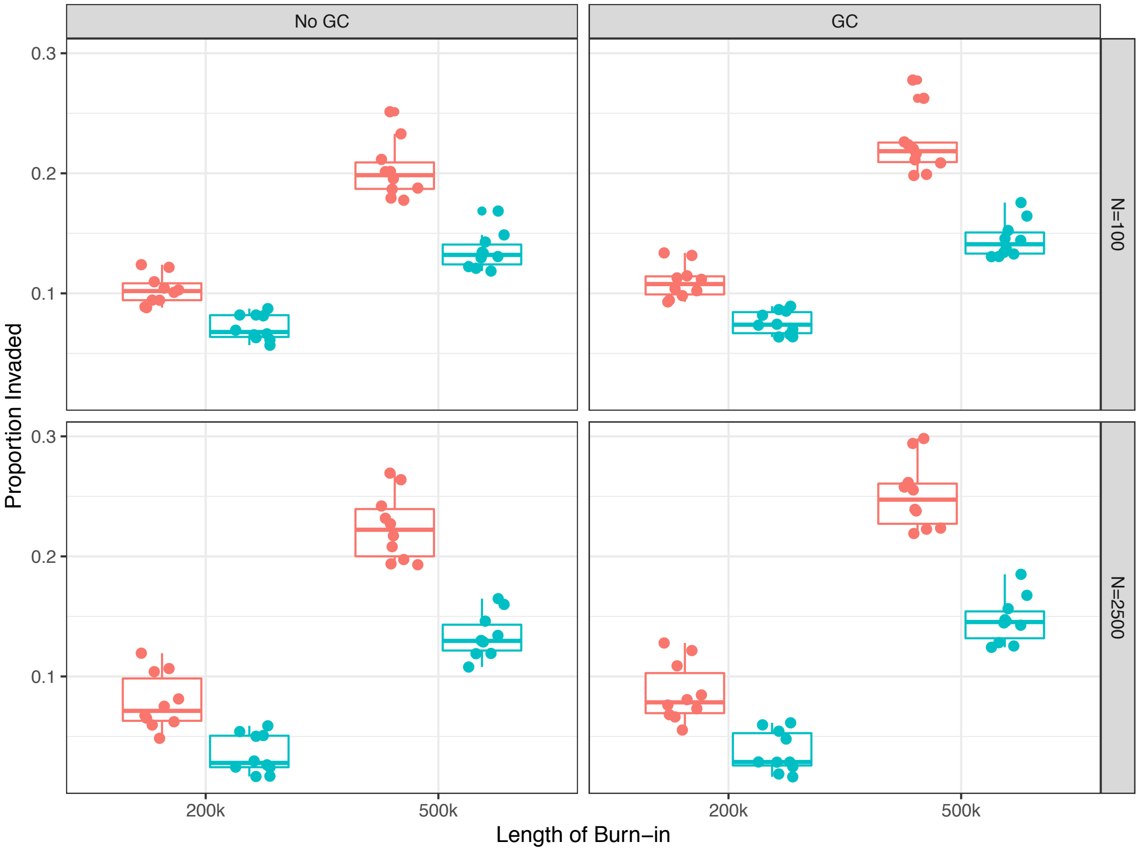


Figure S4. Dominance decreases invasion probability. Probability of invasion is shown for all parameter sets. Facets indicate N (100 or 2,500), and gene conversion (GC; absent or present). Colors indicate the dominance coefficient (0-red, 0.1-blue). Each point is a different burn-in.

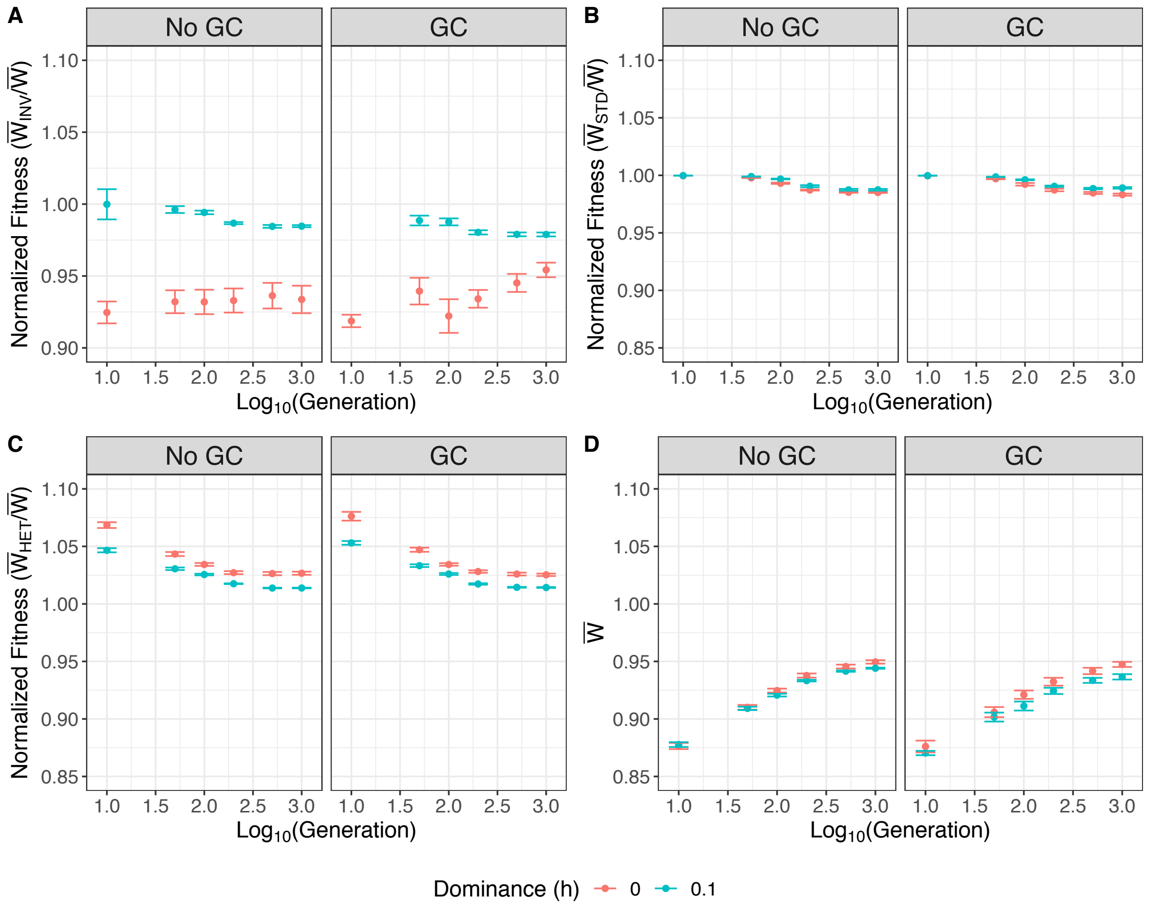


Figure S5. Changes in fitness within the first 1,000 generations after invasion. Shown are normalized fitnesses for A.) the inversion homokaryotype, B.) the standard homokaryotype, C.) the heterokaryotyope, and D.) the mean fitness of the entire population. Normalized fitness for each generation is calculated as an average per burn-in per parameter set. Error bars represent standard error between burn-ins. Facets indicate gene conversion (GC; absent or present) and colors indicate the dominance coefficient (0-red, 0.1-blue). N=2,500 and burn-in length = 500,000 for all graphs.

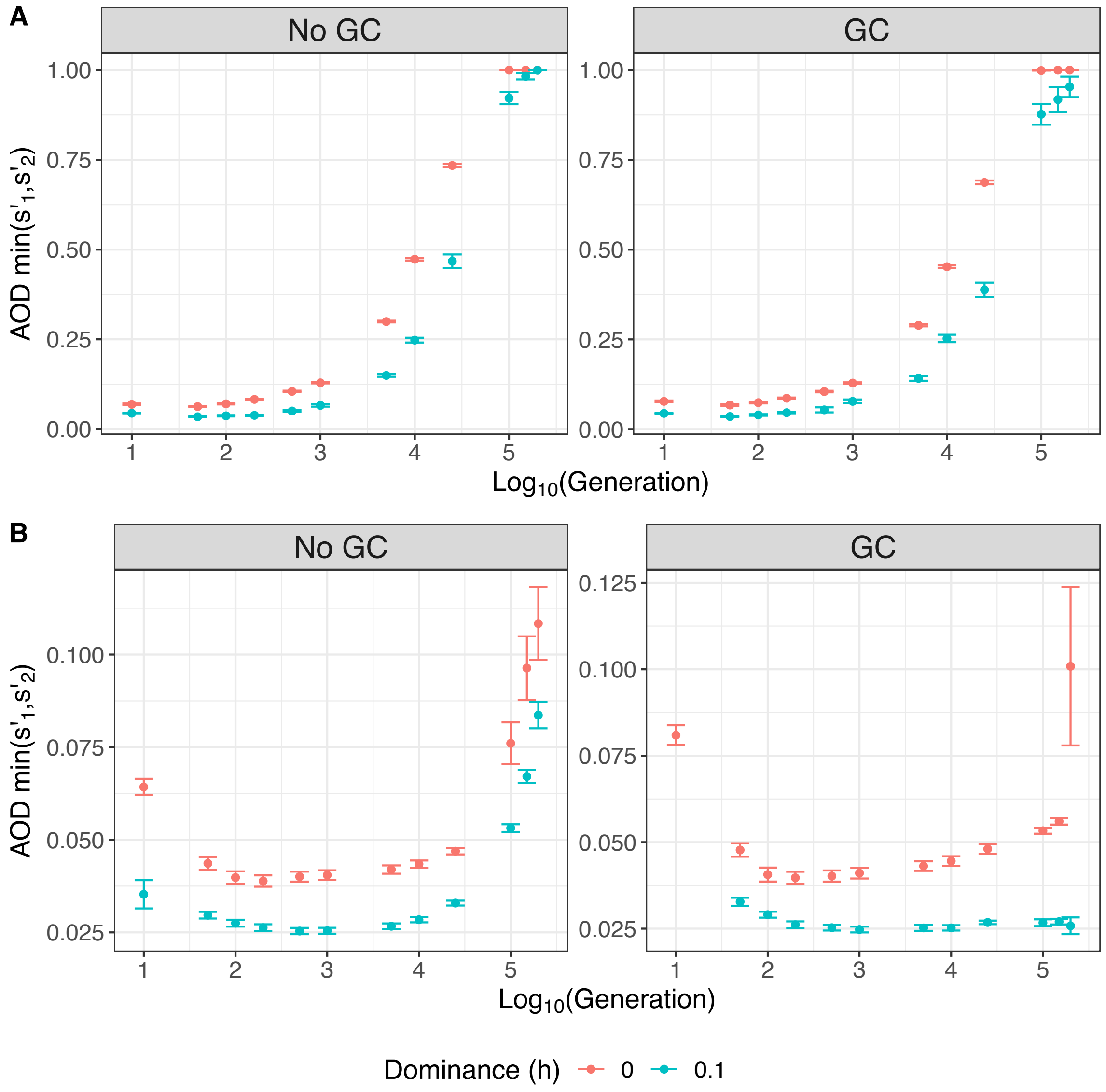


Figure S6. The evolution of associative overdominance (AOD) under different parameters. Shown is the evolution of AOD over time for simulations where the polymorphism lasted 200,000 generations A.) N=100, and B.) N=2,500. AOD for each generation was calculated as an average per burn-in per parameter set. Error bars represent standard error between burn-ins. Colors indicate the dominance coefficient (0-red, 0.1-blue) and facets indicate the level of gene conversion (absent or present). Burn-in length=500,000 for all graphs. Note the change in y-axis scale.

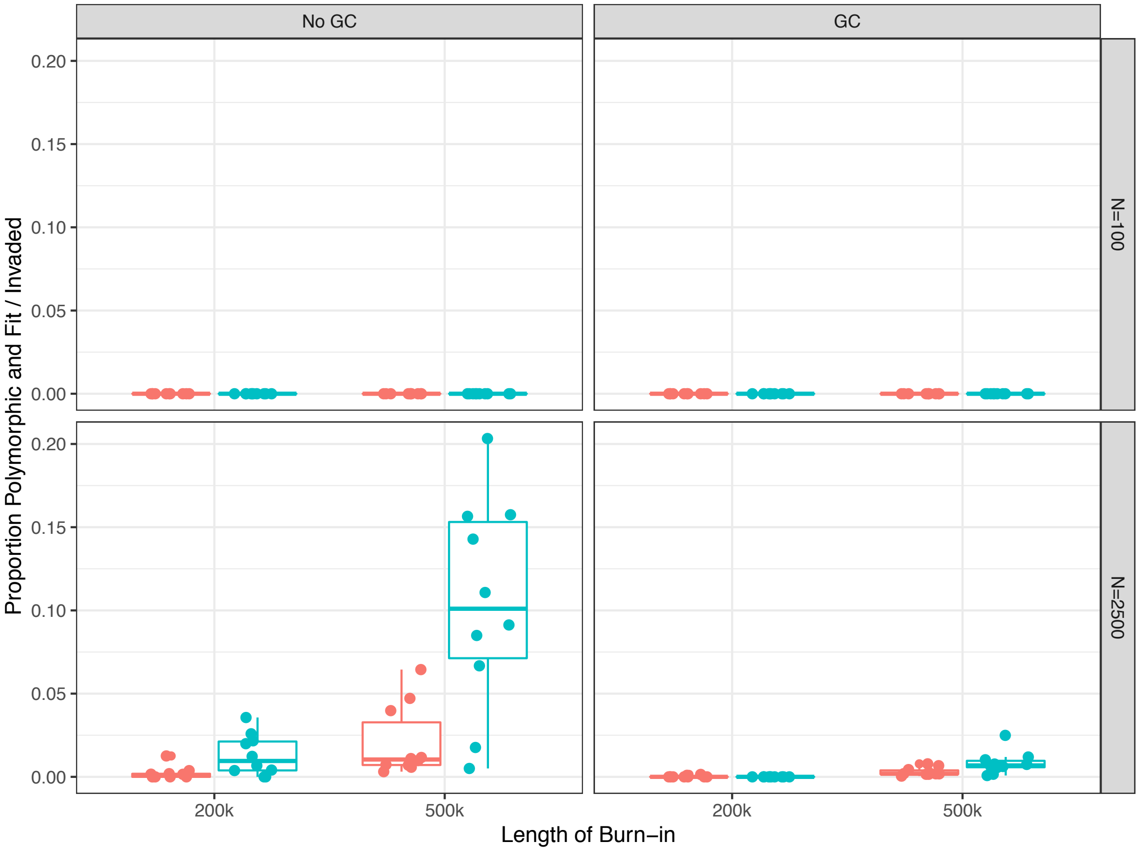


Figure S7. Polymorphism of the inversion with fit homokaryotypes given invasion. Probability that the supergene is polymorphic at the end of the simulation and both homokarotypes are fit is shown for all parameter sets. acets indicate N (100 or 2,500), and gene conversion (GC; absent or present). Colors indicate the dominance coefficient (h; 0-red, 0.1-blue). Each point represents a different burn-in.

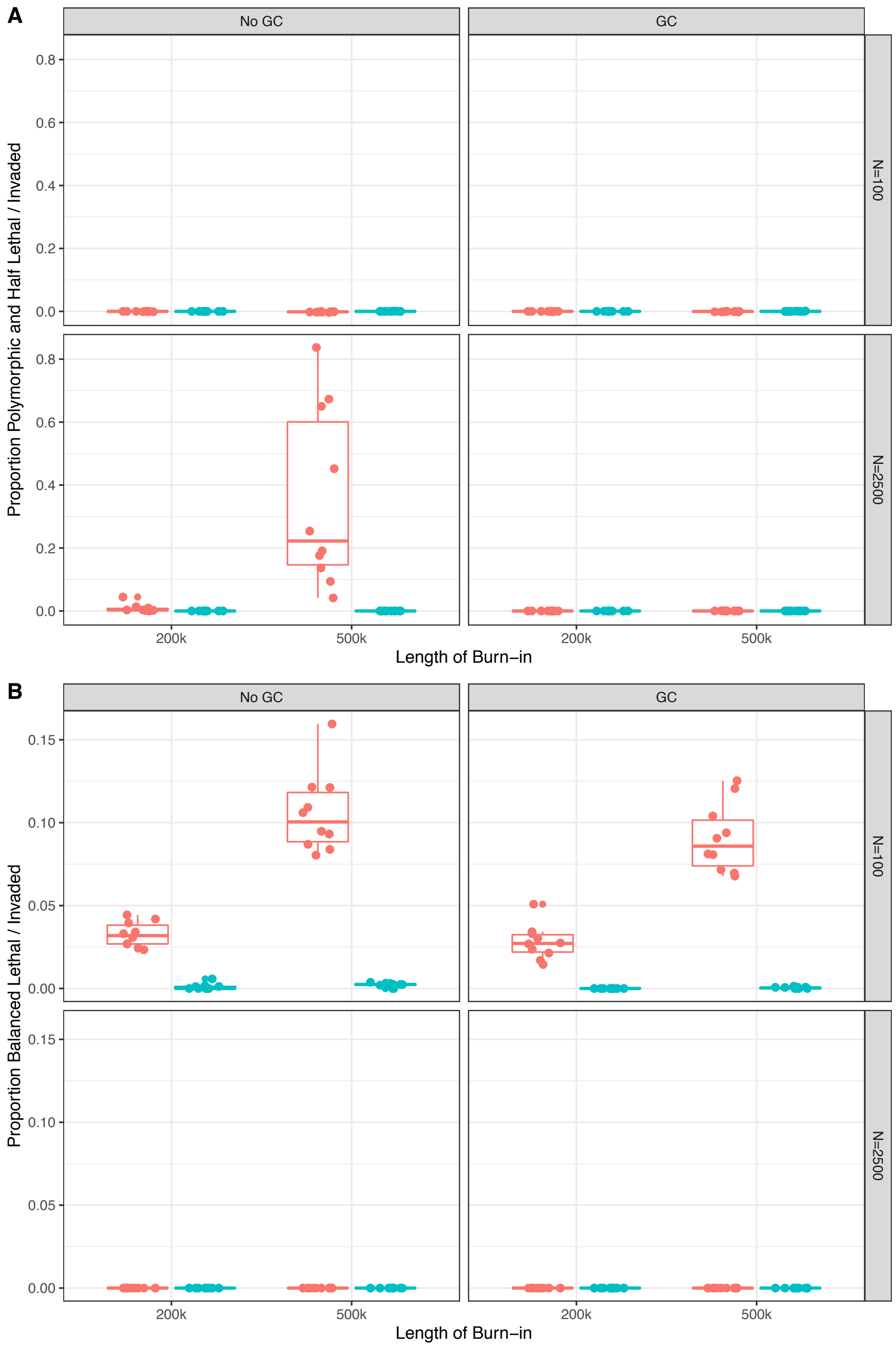


Figure S8. A. Probability of a half lethal system given invasion. Probability that the supergene is polymorphic at the end of the simulation and one homokarotypes is lethal is shown for all parameter sets. B. Balanced lethal outcome given invasion. Probability that the supergene has eveoved into a balanced lethal system after invasion. For both plots facets indicate N (100 or 2,500), and gene conversion (GC; absent or present). Colors indicate the dominance coefficient (h; 0-red, 0.1-blue). Each point represents a different burn-in.

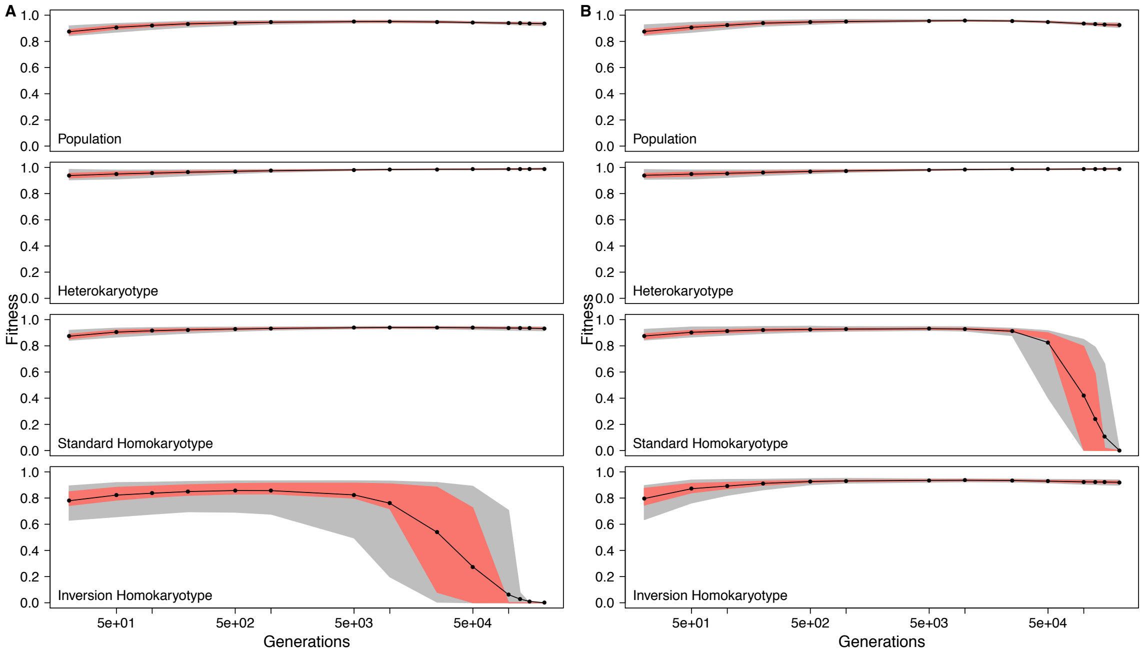


Figure S9. Evolution of karyotype fitnesses in a half lethal system. Fitness of the population, relative fitness of the heterokaryotype, relative fitness of the inversion homokaryotype, and relative fitness of the standard homokaryotype are shown for cases where (A) The inversion homokaryotype becomes lethal, (B) The standard homokarotype becomes lethal. Time is always shown on a log scale, actual time points are: 10, 50, 100, 200, 500, 1000, 5000, 10000, 25000, 50000, 100000, 125000, 150000, and 200000 generations. For all data shown N=2,500, gene conversion is absent and all mutations are fully recessive (h=0).
