## Appendix for "Supergene degeneration opposes polymorphism: The curious case of balanced lethals"

We consider a one-locus model, with overdominance in discrete time. This single, locus $A$, is diallelic, with the fitness of the respective genotype $AA$, $Aa$ and aa given by $w_{AA}$, $w_{Aa}$ and $w_{aa}$. The dynamics of this locus are given by:

$$p_{t+1} = \frac{w_{AA}\ p_t^2 + p_t(1-p_t)\ w_{Aa}}{w_{AA}\ p_t^2 + p_t(1-p_t)\ w_{Aa} + (1-p_t)^2\ w_{aa}} \quad (1)$$

The equilibria of such system are:

$$p = 0;\ p = 1;\ p = \frac{w_{Aa} - w_{aa}}{2w_{Aa} - w_{AA} - w_{aa}} \quad (2)$$

Since we are considering a case of a half-lethal system, we assume $w_{AA} = 0$. In that case for the frequency of allele $A$ to remain above 0.1, the following condition must be verified:

$$9(w_{Aa} - w_{aa}) - w_{Aa} > 0 \quad (3)$$

Using the notation introduced in the main text for associative overdominance, we obtain:

$$9(w_{Aa}\ s'_2)\ -\ w_{Aa} > 0 \quad (4)$$

After normalizing this result by the fitness of the heterozygote Aa, we obtain:

$$9\ s'_2 - 1 > 0 \text{ or } s'_2 > 1/9 \quad (5)$$
